# Temporal integration in human auditory cortex is predominantly yoked to absolute time, not structure duration

**DOI:** 10.1101/2024.09.23.614358

**Authors:** Sam V Norman-Haignere, Menoua K. Keshishian, Orrin Devinsky, Werner Doyle, Guy M. McKhann, Catherine A. Schevon, Adeen Flinker, Nima Mesgarani

## Abstract

Sound structures such as phonemes and words have highly variable durations. Thus, there is a fundamental difference between integrating across absolute time (e.g., 100 ms) vs. sound structure (e.g., phonemes). Auditory and cognitive models have traditionally cast neural integration in terms of time and structure, respectively, but the extent to which cortical computations reflect time or structure remains unknown. To answer this question, we rescaled the duration of all speech structures using time stretching/compression and measured integration windows in the human auditory cortex using a new experimental/computational method applied to spatiotemporally precise intracranial recordings. We observed significantly longer integration windows for stretched speech, but this lengthening was very small (∼5%) relative to the change in structure durations, even in non-primary regions strongly implicated in speech-specific processing. These findings demonstrate that time-yoked computations dominate throughout the human auditory cortex, placing important constraints on neurocomputational models of structure processing.

## Introduction

Natural sounds are composed of hierarchically organized structures that span many temporal scales, such as phonemes, syllables, and words in speech^1^ or notes, contours, and melodies in music^2^. Understanding the neural computations of hierarchical temporal integration is thus critical to understanding how people perceive and understand complex sounds such as speech and music^3-6^. Neural computations in the auditory cortex are constrained by time-limited “integration windows”, within which stimuli can alter the neural response and outside of which stimuli have little effect^7^. These integration windows grow substantially as one ascends the auditory hierarchy^7,8^ and are thought to play a central role in organizing hierarchical computation in the auditory system^5,9^.

A major, unanswered question is whether neural integration in the auditory cortex is tied to absolute time (e.g., 100 milliseconds, **time-yoked integration**) or sound structures (e.g., a phoneme, **structure-yoked integration**). Time- and structure-yoked integration reflect fundamentally different computations because sound structures such as phonemes, syllables, and words are highly variable^10,11^ (**Fig 1A**). The duration of a phoneme, for example, can vary by a factor of 6 or more across talkers and utterances (**Fig 1B**). Thus, if the auditory cortex integrated across speech structures such as phonemes or words – or sequences of phonemes or words – then the effective integration time would necessarily vary with the duration of those structures.

**Figure 1.**
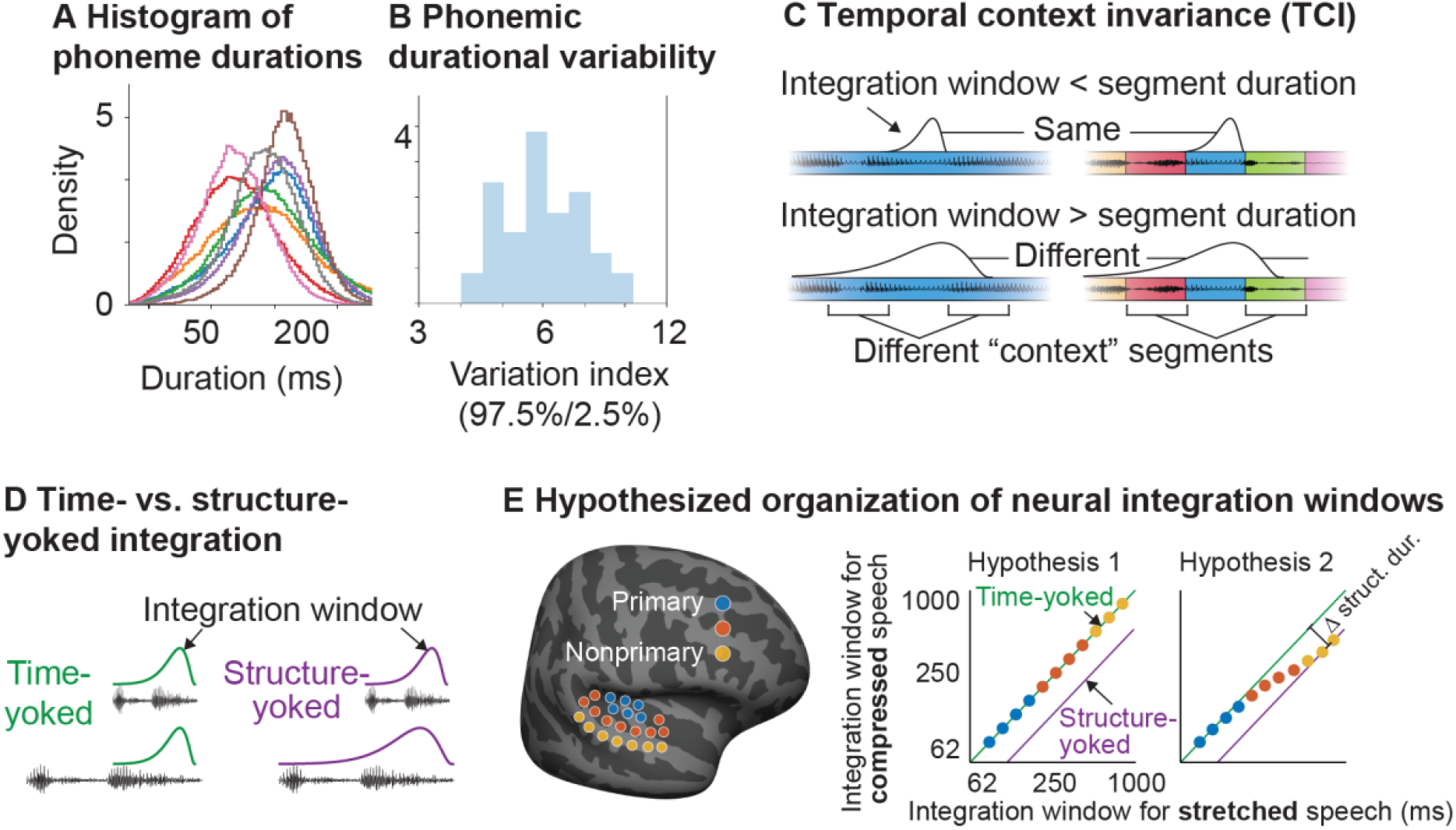
Distinguishing time- and structure-yoked neural integration. **A**, Histogram of phoneme durations across a large speech corpus (LibriSpeech; alignments computed using the Montreal Forced Aligner), illustrating the substantial durational variability of speech structures. Each line corresponds to a single phoneme. **B**, Histogram of the durational variability across all phonemes (median value: 6.0). Durational variability was measured by computing the central 95% interval from the duration histogram for each phoneme (panel A) and then computing a ratio between this interval’s upper and lower bound. This figure plots a histogram of this ratio across all phonemes. **C**, Schematic illustration of the temporal context invariance (TCI) paradigm used to measure integration windows. In this paradigm, each stimulus segment (here speech) is presented in two different “contexts”, where context is simply defined as the stimuli that surround a segment. This panel illustrates two contexts, one where a segment is part of a longer segment and surrounded by its natural context (left) and one where a segment is surrounded by randomly selected alternative segments of the same duration (right) (concatenated using crossfading). If the integration window is less than the segment duration (top), there will be a moment when it is contained within the segment and invariant to the surrounding context. If the integration window is bigger than the segment duration (right), the context can always alter the response. The TCI paradigm estimates the smallest segment duration needed for a context-invariant response. **D**, Schematic illustration of the effect of time compression and stretching on a time-yoked (left, green) and structure-yoked (right, purple) integration window. Compression and stretching rescale the duration of all speech structures and should therefore compress/stretch the integration window if it reflects structure and not time. **E**, Schematic illustration of two hypotheses tested in this study. Each circle plots the hypothesized integration window (logarithmic scale) for stretched (x-axis) and compressed (y-axis) speech for a single electrode with color indicating the electrode’s anatomical location. Time-yoked integration windows will be invariant to stretching/compression (green line) while structure-yoked windows will scale with the magnitude of stretching/compression leading to a shift on a logarithmic scale (purple line). The left panel (Hypothesis 1) illustrates the predicted organization if neural integration windows increased from primary to non-primary regions but remained time-yoked throughout auditory cortex. The right panel (Hypothesis 2) illustrates the predicted organization if there was a transition from time-to structure-yoked integration between primary and non-primary regions.

Currently, little is known about the extent to which neural integration in the auditory cortex reflects time or structure. Auditory models have typically assumed that neural integration is tied to absolute cortical timescales^8,12-15^, for example, by modeling cortical responses as integrating spectral energy within a fixed spectrotemporal receptive field^16,17^ (STRF). In contrast, cognitive and psycholinguistic models have often assumed that information integration is tied to abstract structures such as phonemes or words^4,18-25^. Distinguishing between time- and structure-yoked integration is therefore important for relating auditory and cognitive models, building more accurate neurocomputational models of auditory processing, and interpreting findings from the prior literature.

Distinguishing time- and structure-yoked integration has been difficult in part due to methodological challenges. Temporal receptive field models have been used to characterize selectivity for many different acoustic features (e.g., STRFs) and speech structures^20,25-30^. Temporal receptive field models, however, implicitly assume time-yoked integration, even when applied to sound structures (e.g., phoneme labels), require making strong assumptions about the particular features or structures that underlie a neural response, and cannot account for nonlinear computations, which are prominent in the auditory cortex^31-34^ and likely critical to structure-yoked computations^35^. In addition, standard human neuroimaging methods have poor spatial (e.g., EEG) or temporal (e.g., fMRI) resolution, which is critical for measuring and mapping integration windows in the human auditory cortex.

To address these limitations, we measured integration windows using intracranial recordings from human neurosurgical patients, coupled with a recently developed method for measuring integration windows from nonlinear systems, which does not depend on any assumptions about the features that underlie the neural response or the nature of the stimulus-response mapping (the temporal context invariance or TCI paradigm)^7^ (**Fig 1C**). We then tested whether neural integration windows varied with speech structures by rescaling the duration of all structures using stretching and compression (preserving pitch). Prior studies have measured the degree to which the overall neural response timecourse compresses/stretches with the stimulus^36,37^, but this type of “timecourse rescaling” could be due to changes in the stimulus rather than changes in the neural integration window, as we demonstrate. By contrast, we show that our approach can clearly distinguish time- and structure-yoked integration, including from deep neural network (DNN) models that have complex and nonlinear stimulus-response mappings.

We used uniform stretching/compression in our study, even though it is not entirely natural, because it rescales the duration of all speech structures by the same amount. As a consequence, a structure- yoked integration window should rescale with the magnitude of stretching/compression, irrespective of the particular structures that underlie the window, while a time-yoked window should be invariant to stretching and compression (**Fig 1D**). We tested two primary hypotheses (**Fig 1E**): (1) integration windows increase hierarchically in non-primary regions but remain yoked to absolute time throughout the auditory cortex (2) integration windows become structure-yoked in non-primary auditory cortex.

## Results

### Temporal context invariance effectively distinguishes time-vs. structure-yoked integration

Integration windows are often defined as the time window within which stimuli alter a neural response and outside of which stimuli have little effect^7,38^. This definition is simple and general and applies regardless of the particular features that underlie the response or the nature of the stimulus-response mapping (e.g., linear vs. nonlinear). To estimate the integration window, we measure responses to sound segments surrounded by different “context” segments^7^ (**Fig 1C**). While context has many meanings^39^, here we operationally define context as the stimuli that surround a segment. If the integration window is less than the segment duration, there will be a moment when it is fully contained within each segment and thus unaffected by the surrounding context. In contrast, if the integration window is larger than the segment duration, the surrounding segments can alter the response. We can thus estimate the integration window as the smallest segment duration yielding a context-invariant response. Our stimuli consist of segment sequences presented in a pseudorandom order with shorter segments being excerpted from longer segments. As a consequence, we can compare contexts where a segment is part of a longer segment and thus surrounded by its natural context (**Fig 1C**, left) with contexts where a segment is surrounded by random other segments (**Fig 1C**, right).

We assessed context invariance via the “cross-context correlation”^7^ (**Fig 2A**). We aligned and compiled the response timecourses surrounding all segments as a [segment x time] matrix, separately for each context. We then correlated the corresponding columns across the segment-aligned matrices for each context. Prior to segment onset, the cross-context correlation should be approximately 0, since the integration window must overlap the preceding context segments, which are independent across contexts. As the integration window begins to overlap the shared central segments, the cross-context correlation will rise, and if the window is less than the segment duration there will be a lag when the response is the same across contexts, yielding a correlation of 1, modulo noise.

**Figure 2.**
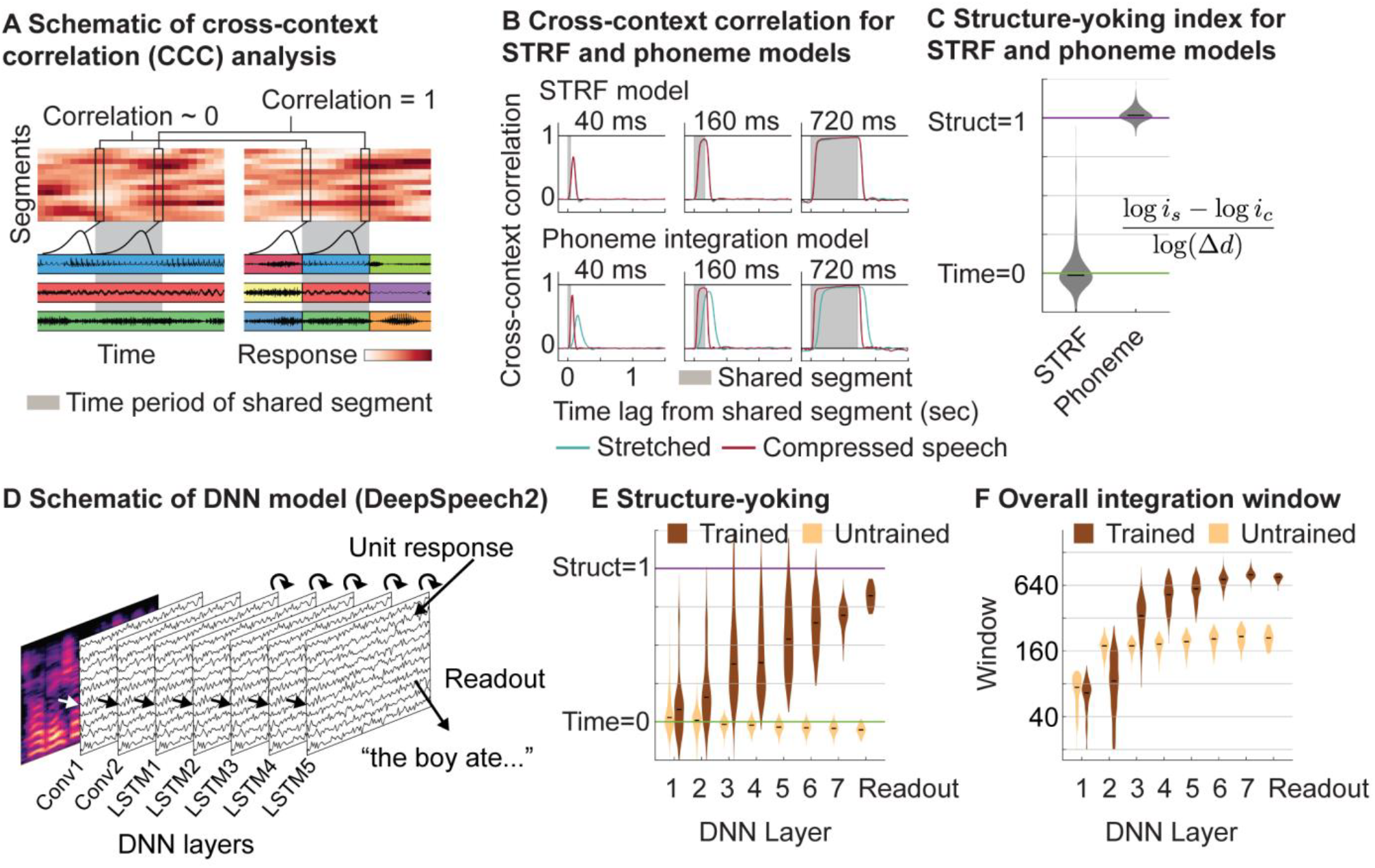
Validating approach using computational models. **A**, Schematic of the cross-context correlation analysis used to measure context invariance for one segment duration. The response time courses to all segments are organized as a [segment x time] matrix, separately for each of the two contexts. Each row contains the response time course to a different segment, aligned to segment onset. The central segments are the same across contexts, but the surrounding segments differ. The grey region highlights the time window when the shared segments are present. To determine whether the response is context invariant, we correlate corresponding columns across matrices from different contexts (“cross-context correlation”), illustrated by the linked columnar boxes. For each box, we plot a schematic of the integration window at that moment in time. At the start of the shared segments (first box pair), the integration window will fall on the preceding context segments, which are random, yielding a cross-context correlation near zero. If the integration window is less than the segment duration, there will be a lag when the integration window is fully contained within the segment, yielding a context-invariant response and correlation of 1 (second box pair). **B**, The cross-context correlation for stretched and compressed speech from two computational models: a model that integrates spectrotemporal energy within a time-yoked window (STRF, top) and a model that integrates phonemes within a structure-yoked window (phoneme integration, bottom). **C**, Structure-yoking index measuring the change in integration windows between stretched and compressed speech on a logarithmic scale (log *i*_*s*_ − log *i*_*c*_), relative to the change in structure durations between stretched and compressed speech (log Δd). The index provides a graded metric measuring the extent of time-(index=0) vs. structure-yoked (index=1) integration. This panel plots the distribution of structure-yoking indices for STRF and phoneme integration models. **D**, Schematic of a deep neural network (DNN) speech recognition model (DeepSpeech2) used to assess whether the TCI paradigm can distinguish time- and structure-yoked integration in a complex, multilayer, nonlinear model. The DNN consists of 8 layers (two convolutional, five recurrent layers, one linear readout), trained to transcribe text from a speech mel-spectrogram without any stretching or compression. Each layer is composed of model units with a response timecourse to sound. Integration windows were estimated by applying the TCI paradigm to the unit timecourses. **E**, Distribution of structure-yoking indices (violin plots) across all units from each layer for both a trained and untrained DNN. **F**, Distribution of overall integration windows, averaged across stretched and compressed speech, for all model units from each layer for both a trained and untrained DNN.

We tested whether we could use our paradigm to distinguish time- and structure-yoked integration by applying the cross-context correlation analysis to responses from computational models. We measured the cross-context correlation for stretched and compressed speech from a spectrotemporal receptive field (STRF) model that linearly integrated spectrotemporal energy within a time-yoked window and a phoneme integration model that linearly integrated phonemic labels within a structure-yoked window whose temporal extent varied with the magnitude of stretching/compression. Both models were fit to cortical responses to natural speech, measured using intracranial recordings, to make them more neurobiologically accurate. We used a stretching/compression factor of 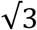 in both our model simulations and neural experiments, because this factor maintains high intelligibility^36,40^, while still producing a relatively large, 3-fold difference in structure durations between compressed and stretched speech.

For both models, the cross-context correlation rose and fell at segment onset/offset and increased with segment duration, as predicted. Notably, the STRF cross-context correlation was nearly identical for stretched and compressed speech, consistent with a time-yoked integration window (**Fig 2B**, top panel), while the phoneme cross-context correlation was lower and more delayed for stretched speech (**Fig 2B**, bottom panel), consistent with a longer integration window. We quantified these effects by computing a structure-yoking index that varied between 0 (time-yoked) and 1 (structure-yoked), calculated by dividing the change in integration windows on a logarithmic scale by the three-fold change in structure duration between stretched and compressed speech (**Fig 2C**). Integration windows were computed as the smallest segment duration needed to achieve a threshold invariance level, as measured by the peak cross-context correlation across lags (threshold set to 0.75; results were robust to the threshold value). We consistently observed structure-yoking values near 0 for our STRF model responses and structure-yoking values near 1 for our phoneme integration models, verifying that we can distinguish time- and structure-yoked integration from ground truth models.

We next examined whether we could detect a transition from time-to structure-yoked integration from a deep neural network (DNN) model (DeepSpeech2) that had been trained to recognize speech structure from sound (trained to transcribe speech from a mel-spectrogram using a connection-temporal-classification loss). DNNs trained on challenging tasks have been shown to learn nonlinear representations that are predictive of cortical responses and replicate important aspects of hierarchical cortical organization, and thus provide a useful testbed for evaluating new methods and generating hypotheses for neural experiments. Moreover, unlike our STRF and phoneme models, the stimulus-response mapping in these models is complex and nonlinear and therefore provides a more challenging setting in which to measure neural integration windows and thus stronger test of our method. We measured integration windows using the same procedure just described but applied to the response of each unit from each layer of the trained DNN model. Importantly, the DNN model was only ever trained on natural speech and thus any structure-yoking present in the model must have been learned solely from the structural variability of natural speech.

This analysis revealed a clear transition from time-to structure-yoked integration across network layers (**Fig 2D**). This transition was completely absent from an untrained model, demonstrating that it was learned from the structural variability of natural speech. The overall integration time, averaged across stretched and compressed speech, also increased substantially across network layers for the trained model (**Fig 2E**), an effect that was also primarily due to training (the increase from layer 1 to layer 2 is present in the random network because of striding in the architecture). These results demonstrate that we can distinguish time-and structure-yoked integration, as well as transitions between the two using our approach, including from complex, nonlinear models that have only ever been trained on natural speech. These results also help motivate our second hypothesis (**Fig 1E**, right panel), since prior studies have found that later network layers better predict later-stage regions of the human auditory cortex^41^, suggesting that there may be a transition from short, time-yoked integration to long, structure-yoked integration across the auditory hierarchy.

### Neural integration throughout human auditory cortex is predominantly time-yoked

We next sought to test whether integration windows in the human auditory cortex reflect time-or structure-yoked integration. We measured cortical responses to speech segments (37, 111, 333, 1000, 3000 ms) from an engaging spoken story (from the Moth Radio Hour), following time compression or stretching (preserving pitch). Our paradigm was designed to characterize sub-second integration windows within the auditory cortex and it was not possible to characterize multi-second integration windows beyond the auditory cortex due to the small number of segments at these longer timescales. All analyses were performed on the broadband gamma power response of each electrode (70-140 Hz). We focus on broadband gamma because it provides a robust measure of local electrocortical activity^42,43^ and can be extracted using filters with narrow integration windows (∼19 ms), which we have shown have a negligible effect on the estimated integration window^7^. In contrast, low-frequency, phase-locked activity requires long-integration filters that substantially bias the measured integration window^7^.

The cross-context correlation for stretched and compressed speech is shown for two example electrodes, one from a primary region overlapping right Heschl’s gyrus (HG) and one from a non-primary region overlapping the right superior temporal gyrus (STG) (**Fig 3A**). Because neural data is noisy (unlike model responses), we plot a noise ceiling for each electrode which measures the cross-context correlation when the context is the same using repeated presentations of the same segment sequence. The STG electrode required longer segment durations and lags for the cross-context correlation to reach the noise ceiling, indicating a longer integration window. But notably, the cross-context correlation was similar for stretched and compressed speech for both the HG and STG electrode, suggesting a window predominantly yoked to absolute time.

**Figure 3.**
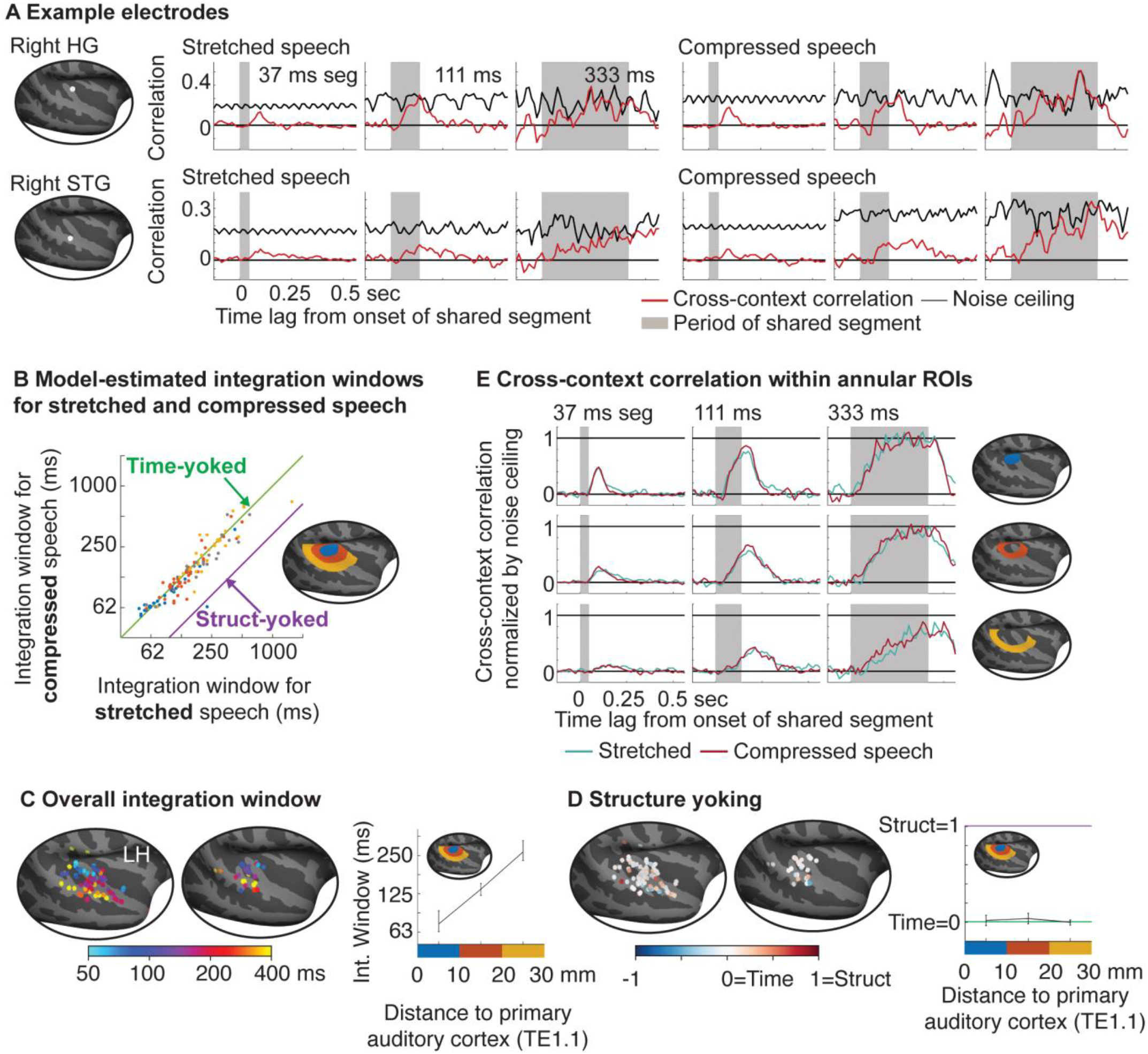
Neural integration in human auditory cortex is predominantly time-yoked. **A**, Cross-context correlation (red line) and noise ceiling (black line) for stretched (left) and compressed (right) speech from two example electrodes overlapping right Heschl’s gyrus (HG) (top) and right superior temporal gyrus (STG) (bottom). **B**, Integration windows for stretched (x-axis) and compressed (y-axis) speech for all sound-responsive electrodes. Green and purple lines indicate what would be predicted from a time- and structure-yoked integration window. Electrodes are colored based on annular regions-of-interest (ROI) that reflect the distance of each electrode to primary auditory cortex (inset; gray colors reflect electrodes outside the annular ROIs). **C**, Map plotting the average integration window across stretched and compressed speech. Right panel plots the median integration window within annular ROIs. Error bars plot one standard error of the bootstrapped sampling distribution across subjects and electrodes. **D**, Map of structure yoking. Right panel plots the median structure-yoking index within annular ROIs. **E**, Cross-context correlation (normalized by noise ceiling) for stretched and compressed speech averaged across all electrodes within each annular ROI (top to bottom).

We used a parametric model developed in our prior work that makes it possible to estimate integration windows from noisy neural data by pooling across all lags and segment durations, allowing us to quantify the integration window for each electrode for both stretched and compressed speech (**Fig 3B**; **Fig S1&2**). Electrodes were localized on the cortical surface, which we used to create a map of the overall integration window (**Fig 3C**), averaged across stretched and compressed speech, as well as a map of structure yoking (**Fig 3D**). As in prior work^7,33^, we quantified differences in neural integration related to the cortical hierarchy by binning electrodes into annular ROIs based on their distance to primary auditory cortex (three 10-mm spaced bins) (**Fig 3B-E**). As a simple, model-free summary metric, we also computed the average cross-context correlation across all electrodes within each ROI separately for stretched and compressed speech and normalized the cross-context correlation by the noise ceiling at each lag and segment duration (**Fig 3E**). For our annular ROI analyses, we pooled across hemispheres because integration windows were similar between the right and left hemispheres both in this study (**Fig 3D&E**) and in our prior work^7^, and because we had a small sample size in the right hemisphere (33 electrodes).

We found that the overall integration window, averaged across stretched and compressed speech, increased substantially across the cortical hierarchy with an approximately three-fold increase from primary to non-primary regions (**Fig 3B, C**) (median integration for the annular ROIs: 80, 130, and 272 ms for annular ROIs 1; *F*_1,9.49_ = 20.65, *p < 0*.*01, β* = 0.088 *octaves/mm, CI* = /0.050, 0.127], 108 electrodes, 15 subjects, Linear Mixed Effects model). The cross-context correlation showed a lower peak value and a more gradual build-up and fall-off in ROIs further from primary auditory cortex, again consistent with a longer integration time (**Fig 3E**). These findings replicate prior work^7^ showing that the human auditory cortex integrates hierarchically across time with substantially longer integration windows in higher-order regions.

We next investigated structure yoking. Across all electrodes, we observed a significant increase in integration windows for stretched compared with compressed speech (**Fig 3B**) (*F*_1,110_ = 8.58, *p < 0*.*01, β* = 0.097 *octaves, CI* = /0.031, 0.163], 110 electrodes, 15 subjects). The magnitude of this increase, however, was small relative to the three-fold difference in speech structure durations. Specifically, the median change between stretched and compressed speech was only 0.06 octaves, yielding a structure yoking index of 0.04 when compared with the 1.58-octave difference in structure durations between stretched and compressed speech. Notably, structure yoking was similarly weak throughout both primary and non-primary auditory cortex (**Fig 3D**): structure yoking did not increase with distance from primary auditory cortex (*F*_1,7.67_ = 0.36, *p* = 0.57, *β* = ™0.007 Δ*octaves/mm, CI* = /™0.031, 0.017], 108 electrodes, 15 subjects) and did not become stronger in electrodes with longer overall integration times (*F*_1,29.87_ = 0.22, *p* = 0.64, *β* = 0.018 Δ*octaves/octaves, CI* = /™0.060, 0.097], 110 electrodes, 15 subjects), as predicted by Hypothesis 2. The average cross-context correlation was very similar for stretched and compressed speech even in annular ROIs far from primary auditory cortex, again consistent with time-yoked integration (**Fig 3E**). These findings suggest that the auditory cortex integrates hierarchically across time, but that the fundamental unit of integration is absolute time and not structure.

### Timecourse rescaling does not distinguish time- and structure-yoked integration

Prior studies have observed that stretching or compressing speech causes the neural response timecourse to stretch and compress^36,37^. We replicated these prior studies by correlating the response timecourse of each electrode to stretched and compressed speech after time-stretching the neural response to compressed speech (**Fig 4A**) (by the 3-fold difference in speech rates), and then dividing by the maximum possible correlation given the data reliability. Consistent with prior studies, we observed substantial timecourse rescaling values throughout primary and non-primary auditory cortex (**Fig 4B**) (median rescaling value across the three annular ROIs was 0.47, 0.43, and 0.39) (*F*_1,12.40_ = 40.48, *p <* 0.001, *β* = 0.063, *CI* = /0.044, 0.083], 131 electrodes, 15 subjects).

**Figure 4.**
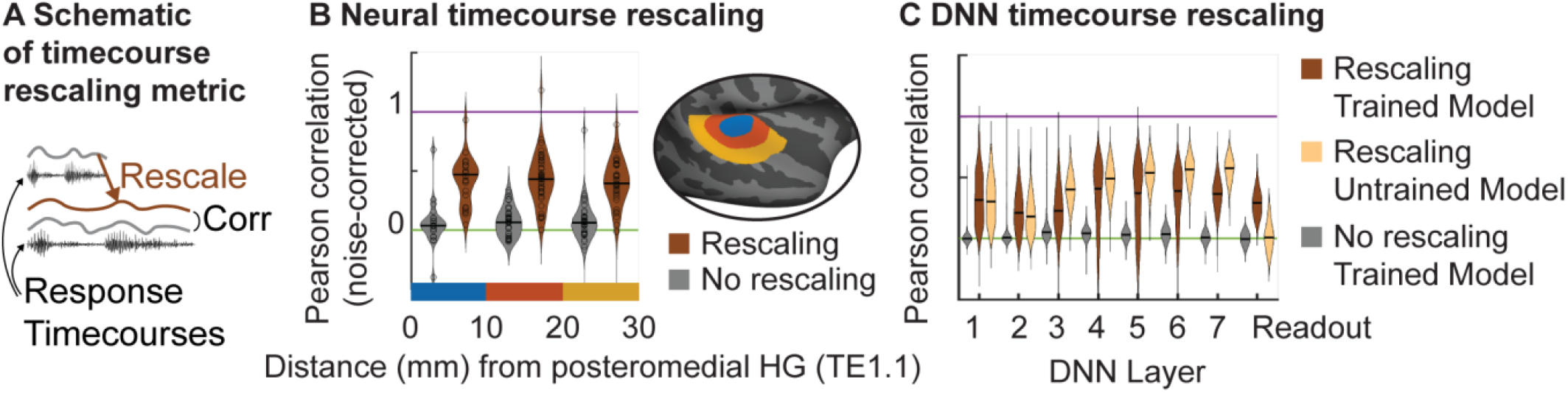
Timecourse rescaling does not distinguish time- and structure-yoked integration. **A**, The timecourse for compressed speech was rescaled (stretched) using resampling and then correlated with the neural response timecourse for stretched speech. **B**, The correlation between stretched and compressed speech with and without rescaling (noise-corrected by response reliability). Electrodes were grouped into annular ROIs and violin plots show the distribution of the timecourse rescaling metric across all electrodes within each ROI. **C**, Timecourse rescaling for different layers of a trained and untrained DNN model, as well as baseline correlations without rescaling (for the trained model).

Neural timecourse rescaling might naively be interpreted as reflecting a rescaled (structure-yoked) integration window, but might alternatively reflect a rescaled stimulus with a fixed (time-yoked) integration window. To address this question, we examined timecourse rescaling in our DNN model, which showed a clear, training-dependent change in structure-yoked integration across its layers. We found that all DNN layers showed substantial timecourse rescaling, even the earliest layers, and that rescaling was similar for trained and untrained models (**Fig 4C**). These findings contrast sharply with the results from our TCI analysis (**Fig 2E**) and suggest that timecourse rescaling provides a poor metric of structure-yoked integration, being driven to a large extent by stimulus rescaling rather than integration window rescaling. This finding underscores the significance of our approach, which can cleanly isolate time- and structure-yoked integration from complex, nonlinear responses.

## Discussion

We tested whether the human auditory cortex integrates information across absolute time or sound structure in speech. Leveraging our recently developed TCI method, we developed a novel approach that we demonstrated was effective at distinguishing time- and structure-yoked integration in computational models, including revealing a transition from time- to structure-yoked integration in nonlinear, DNN models trained to recognize structure from natural speech. We then applied our approach to intracranial recordings from human neurosurgical patients, which have high spatiotemporal precision, enabling us to measure integration windows throughout primary and non-primary human auditory cortex. Across all electrodes, we observed a significant increase in integration times for stretched compared with compressed speech. This change, however, was small relative to the difference in structure durations between stretched and compressed speech, even in non-primary regions of the STG strongly implicated in structure processing^6,44,45^. These findings demonstrate that the primary unit of integration throughout the human auditory cortex is absolute time and not structure duration.

### Implications for computational models

Auditory and cognitive neuroscientists have often studied sound processing using distinct sets of models and assumptions. Auditory models typically assume that neurons integrate simple types of acoustic features over fixed temporal windows^8,12-15^ (e.g., STRFs). These models have been successful in explaining neural responses early in the auditory system, but have had greater difficulty in predicting neural responses to natural sounds in higher-order regions of the auditory cortex^31,41^. For example, prior work has revealed prominent selectivity for speech and music that cannot be explained by spectrotemporal receptive fields^6,33,45^, as well as tuning for speech-specific structures such as phonemes and phonotactics^20,26-30^ (STG).

Cognitive models, by contrast, have often assumed that temporal integration is yoked to abstract structures such as phonemes or words^4,18-25^, often leaving unspecified the acoustic computations used to derive these structures from sound. For example, models of spoken recognition are often cast as a sequential operation applied to phonemes^19,46^, and features inspired by these models have shown promise in predicting cortical responses in non-primary auditory cortex^20,47,48^. Language models, which integrate semantic and syntactic information across words (or “word pieces”) have also shown strong predictive power, including in non-primary regions of the auditory cortex such as the STG^21-24^. Collectively, these observations suggest that there might be a transition from time-yoked integration of acoustic features in primary regions to structure-yoked integration of abstract structures such as phonemes or words in non-primary regions.

Our findings are inconsistent with this hypothesis since we find that neural integration is predominantly yoked to absolute time throughout both primary and non-primary auditory cortex with little change in structure yoking between primary and non-primary regions. These findings do not contradict prior findings that non-primary auditory cortex represents speech- and music-specific structure, but do suggest that the underlying computations are not explicitly aligned to sound structures as in many cognitive and psycholinguistic models. For example, phonotactic models that compute measures of phoneme probability using a sequence of phonemes^19,46^, implicitly assume that the neural integration window is yoked to phonemes and thus varies with phoneme duration. Similarly, language models that integrate across words and word pieces implicitly assume that the integration window varies with word duration^21-24^. Our findings indicate that these types of structure-yoked computations are weak in the auditory cortex, which has important implications for models of neural coding. For example, a corollary of our finding is that the amount of information analyzed by the auditory cortex will be inversely proportional to the rate at which that information is presented since compressing speech increases the information present within the integration window, and stretching speech decreases the information within the integration window. Similarly, the number of phonemes and words that the auditory cortex effectively analyzes will be less when those phonemes and words have longer durations.

How can people recognize speech structures using time-yoked integration windows, given their large durational variability? One possibility is that integration windows in the auditory cortex are sufficiently long to achieve recognition of the relevant sound structures, even if they are yoked to absolute time. Non-primary regions integrate across hundreds of milliseconds and do not become fully invariant to context even at segment durations of 333 ms (**Fig 3E**), and thus will integrate across many phonemes even if those phonemes have long durations. This may be analogous to higher-order regions of visual cortex that have large spatial receptive fields, sufficient to recognize objects across many spatial scales^49,50^. Structure-yoked computations may also be instantiated in downstream regions, such as the superior temporal sulcus or frontal cortex that integrate across longer, multisecond timescales^3,4,51^, either by enhancing weak structure-yoked computations already present in the auditory cortex or by explicitly aligning their computations to speech structures and structural boundaries^19,52^.

Our findings are broadly consistent with anatomical models that posit a hierarchy of intrinsic timescales, driven by hierarchically organized anatomical and recurrent connections^53,54^. Since intrinsic timescales are stimulus-independent they are by definition time-yoked and will not vary with structure duration. Stimulus-independent dynamics can nonetheless influence the integration window by controlling the rate at which sensory information is accumulated and forgotten within neural circuits. Several recent neurocognitive models have posited that neural integration in the cortex is yoked to “event boundaries” in speech, such as the boundary between words, sentences, or longer narrative structures^55,56^. Our results rule out simple models of event-based integration within the auditory cortex, such as models where the integration window grows based on the distance to a structural boundary^57^, since the distance scales with the magnitude of stretching and compression.

### Relationship to prior methods

Many methods have been developed to study the temporal characteristics of neural responses, but in most cases, these methods do not provide a direct estimate of the integration window. For example, many prior studies have measured the autocorrelation of the neural response in the absence of or after removing stimulus-driven responses, as a way to assess intrinsic timescales^53,58^. Intrinsic timescales are thought to in part reflect network dynamics such as the integration time of a neural population with respect to its synaptic inputs^54^. The neural integration window, as defined here, specifies the integration time of a neuron with respect to the stimulus, and includes the cumulative effects of all network dynamics on stimulus processing, both feedforward, feedback, and recurrent.

Many prior studies have measured temporal properties of the neural response timecourse, such as frequency characteristics of the response^59,60^ or the degree to which the neural response tracks or phase locks to the stimulus^61-63^. These methods while useful, do not provide a direct measure of the neural integration window in part because they are influenced by both the stimulus and neural response characteristics. For example, stretching the stimulus will tend to stretch the neural response (changing the frequency spectrum as well) even in the absence of a change in neural integration, which explains why we observed strong timecourse rescaling for neural responses with time-yoked integration windows. This observation underscores the utility of having a method like the TCI paradigm that can directly estimate the neural integration window, independent of stimulus characteristics, in order to distinguish time- and structure-yoked integration.

Temporal receptive models can be conceptualized as estimating the integration window of a best-fitting linear system, for example, between a spectrogram-like representation and the neural response in the case of a STRF^8,16,17^. However, as our DNN analyses show, nonlinear processing is likely critical to instantiating structure-yoked computations, and it is therefore important to be able to measure the integration window of a nonlinear system to distinguish between time- and structure-yoked integration. Temporal receptive field models have also been applied to investigate selectivity for speech structures, such as phonemes or phonotactic features^20,26-30^, which implicitly assumes time-yoked integration of these structures. By demonstrating strong, time-yoked integration throughout the auditory cortex, our findings provide some justification for this assumption.

Several studies have reported neural responses that change at structural event boundaries in speech or music^56,64^, including in the auditory cortex, which has motivated event-based integration models^55^. However, many features of sound change at structural boundaries, which could produce a neural response at the boundary even if the integration window does not change at the boundary. These prior findings are thus not inconsistent with our study, and more research is needed to determine whether there are any event-based changes in integration within the auditory cortex.

Many studies have measured selectivity for longer-term temporal structure by comparing intact and temporally scrambled sounds^3,4,6,51^. These scrambling metrics however do not provide a direct measure of the integration window, since many regions of the auditory cortex (e.g., primary auditory cortex) show no effect of scrambling even at very short timescales^6^ (e.g., 30 milliseconds) despite having a meaningful integration window at these durations^7^. Here, we were able to identify a wide range of integration times throughout primary and non-primary auditory cortex by combining our TCI method with the spatiotemporal precision of intracranial recordings. We were able to detect a small change in integration windows between stretched and compressed speech across auditory cortex, while simultaneously showing that this change was small relative to the magnitude of stretching and compression and similar between primary and non-primary regions. Our methods were thus critical to distinguishing between time- and structure-yoked integration and provide clear evidence that integration windows in the auditory cortex are predominantly time-yoked.

### Methodological choices, limitations, and future directions

We chose to use uniform stretching and compression so that all speech structures would be stretched or compressed by the same amount. This choice was made because speech contains many perceptually relevant structures and we do not know a priori which structures will underlie a particular neural response. Although uniform stretching and compression are not entirely natural, people can still understand speech with moderate amounts of uniform stretching and compression^36,40^, like that used in this study, and thus whatever mechanisms are used by the brain to recognize speech must still be operational in this regime. Moreover, we showed that we could use uniform stretching and compression to detect a transition to structure-yoked integration from a nonlinear DNN model that had only ever been trained on natural speech (**Fig 2E**).

We chose to use segments with a fixed duration because this simplified our analyses. Fixed-duration segments contain mixtures of full and partial structures (e.g., half a word), but this property is not problematic for identifying structure-yoked integration, because the same mixtures are present for both stretched and compressed speech. For example, if a structure-yoked population only responded strongly to complete words then the segment duration needed to achieve a strong response would still be three times as long for stretched vs. compressed speech since the duration needed for a given number of complete words to be present would be three times as long for the stretched speech. Our computational models verified that we could detect structure-yoked integration using fixed duration segments.

Our analysis focused on broadband gamma power because it provides a standard measure of local electrocortical activity and because it can be extracted with short-duration filters that have little effect on the measured integration^7^. In contrast, low-frequency phase-locked activity is measured using filters that are long and fixed, which inevitably biases responses toward long, time-yoked integration. Future work could potentially examine lower-frequency, phase-locked activity using spatial filters designed to extract such activity rather than temporal filters^65^.

The large majority of the segments tested in this experiment were less than a second to allow us to measure integration windows from the auditory cortex which has sub-second integration times^7^. Regions beyond the auditory cortex in the superior temporal sulcus and frontal cortex show longer, multi-second integration windows^3,4,51^. Future work could examine whether these higher-order regions show time- or structure-yoked integration windows by testing a larger number of long-duration segments, although this is experimentally challenging because longer-duration segments inevitably require longer experiment times.

## Methods

### Measuring durational variability of speech phonemes

To illustrate the variability of speech structures, we measured the durational variability of phonemes in the LibriSpeech corpus^66^. Phoneme alignments were computed using the Montreal Forced Aligner^67^. For each phoneme, we then measured the distribution of durations across all speakers and utterances in the corpus. We then calculated the central 95% interval of this distribution and measured the ratio between the upper and lower boundaries of this interval as a measure of durational variability.

### Cross-context correlation analysis

We used the “cross-context correlation” to estimate context invariance from both computational models and neural data. In this analysis, we first compile the response timecourses to all segments of a given duration in a [segment x time] matrix (**Fig 2A**). Each row contains the response time course surrounding a single segment, aligned to segment onset. Different rows thus correspond to different segments, and different columns correspond to different lags relative to segment onset. We compute a separate matrix for each of the two contexts being compared. The central segment is the same between contexts, but the surrounding segments differ.

Our goal is to determine whether there is a lag when the response is the same across contexts. We instantiate this idea by correlating corresponding columns across segment-aligned response matrices from different contexts (schematized by the linked columnar boxes in **Fig 2A**). At segment onset (**Fig 2A**, first box pair), the cross-context correlation should be near zero since the integration window must overlap the preceding segments, which are random across contexts. As time progresses, the integration window will start to overlap the shared segment, and the cross-context correlation should increase. If the integration window is less than the segment duration, there will be a lag where the integration window is fully contained within the shared segment, and the response should thus be the same across contexts, yielding a correlation of 1 (**Fig 2A**, second box pair).

Our stimuli enable us to investigate and compare two types of contexts: (1) cases where a segment is a subset of a longer segment and thus surrounded by its natural context (**Fig 1C**, left) and where a segment is surrounded by randomly selected segments of the same duration (**Fig 1C**, right). We computed the cross-context correlation by comparing natural and randomly selected contexts, as well as two different randomly selected contexts (see prior paper^7^ for details), but results were very similar when only comparing natural and random contexts. We note that any response that is selective for naturalistic structure (e.g., a word) will by definition show a difference between natural and random contexts and thus our paradigm will be sensitive to this change.

### Estimating integration windows from computational model responses

We tested whether our approach could distinguish time- vs. structure-yoked integration by applying our analyses to the outputs of computational models. Our methods were very similar to our neural analyses, except that in the case of computational models, the response is noise-free and we are not constrained by experiment time and thus can measure responses to many segments. We measured the cross-context correlation using 30 segment durations with 100, 27-second sequences per duration (segment durations: 20, 40, 60, 80, 120, 160, 200, 240, 280, 320, 400, 480, 560, 640, 720, 800, 880, 960, 1040, 1120, 1200, 1280, 1440, 1600, 1760, 1920, 2080, 2240, 2400, 2560 ms). Segments were excerpted from LibriSpeech following time compression and stretching by a factor of 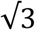 as in the neural experiments (time stretching and compression were implemented using waveform synchronous overlap and add, as implemented by SOX in Python^68^).

To estimate the integration window, we applied our cross-context correlation analysis, which is described both in the main text and in our prior paper^7^ (**Fig 2A**) (comparing natural and random contexts). We then calculated the peak cross-context correlation value for each segment duration as a measure of context invariance. Finally, we interpolated the correlation vs. segment duration curve to determine the smallest segment duration needed to achieve a threshold cross-context correlation value (threshold set to 0.75).

#### Spectrotemporal receptive field model

Following standard practice, our spectrotemporal receptive field (STRF) model was defined by applying a linear transformation to a spectrogram representation of sound (mel spectrogram with torchaudio; sample_rate=16000, n_fft=1280, hop_length=320, f_min=40, f_max=8000, n_mels=128, power=2.0, norm=‘slaney’). To make the STRF model more realistic, the STRFs were fit to approximate human cortical responses to natural speech recorded intracranially from patients. The data and stimuli used to fit the STRF models have been described previously^30^ and consisted of 566 electrode responses from 15 patients. Each patient listened to 30 minutes of speech excerpted from a children’s storybook (*Hank the Cowdog*) and an instructional audio guide (four voice actors; 2 male, 2 female). The STRFs were fit using standard regularized regression with cross-validation used to select the regularization parameter^69^ (10-fold cross-validation). To suppress edge artifacts^70^, a 50 ms half-Hanning window was applied to the beginning and end of each TRF.

#### Phoneme integration model

As a simple model of structure-yoked integration, we instantiated a model that integrated phonemic features within a window whose temporal extent varied inversely with the speech rate. There were 22 phoneme features, each indicating the presence or absence of a single feature (e.g., manner of articulation)^30^. Results were very similar when applied to binary phoneme labels rather than phoneme features. The integration window was determined by fitting the TRF model to the same dataset of responses to natural speech described above, and then stretching or compressing the TRF (via interpolation) by the same stretching and compression factor applied to the speech stimulus.

#### Deep neural network model

We computed integration windows from a popular DNN speech recognition model (DeepSpeech2)^35,71^. The DNN consists of two convolutional layers, five recurrent layers (long-short-term memory cells), and one linear readout layer and was trained to transcribe text from a mel spectrogram using a connection-temporal-classification (CTC) loss (applied to graphemes). The model was trained using 960 hours of speech from the LibriSpeech corpus using standard data augmentation techniques^35^ (background noise, reverberation, and frequency masking) to make recognition more challenging (25 epochs; optimization was implemented in PyTorch using the Adam optimizer). Each layer is defined by a set of unit response time courses and we measured the integration of each unit time course by applying the TCI analysis to its response (in the same manner as all other models). Like most DNN models, DeepSpeech2 is acausal, but this is not problematic for measuring its integration window^35^, because acausality shifts the cross-context correlation to earlier lags and our measure of context invariance (the peak correlation across lags) is invariant to shifts.

### Intracranial recordings from human patients

#### Participants and data collection

Data come from 15 patients undergoing treatment for intractable epilepsy at the NYU Langone Hospital (6 patients) and the Columbia University Medical Center (9 patients) (7 male, 8 female; mean age: 36 years, standard deviation: 13 years). No formal tests were used to determine the sample size, but the number of subjects was larger than in most intracranial studies, which typically test fewer than 10 subjects^5,36^. Electrodes were implanted to localize epileptogenic zones and delineate these zones from eloquent cortical areas before brain resection. NYU patients were implanted with subdural grids, strips, and depth electrodes depending on the clinical needs of the patient. CUMC patients were implanted with depth electrodes. All subjects gave informed written consent to participate in the study, which was approved by the Institutional Review Boards of CUMC and NYU. NYU patients were compensated $20/hour. CUMC patients were not compensated due to IRB prohibition.

#### Stimuli

Segments of speech were excerpted from a recording of a spoken story from the Moth Radio Hour (Tina Zimmerman, *Go In Peace*). The recording was converted to mono, resampled to 20 kHz, and pauses longer than 500 ms were excised. The stimuli were then compressed and stretched while preserving pitch using a high-quality speech vocoder STRAIGHT^72^. There were five, logarithmically spaced segment durations (37 ms, 111 ms, 333 ms, 1000 ms, 3000 ms). Each stimulus was 27 seconds and was composed of a sequence of segments of a single duration. There were two stimuli per segment duration, each with a different ordering of segments. We also tested a 27-second stimulus composed of a single, undivided excerpt. Shorter segments were created by subdividing longer segments. Each stimulus was repeated twice (in one subject, the stimuli were repeated four times). In two participants, we tested an additional segment duration (9 seconds) and used a longer stimulus duration (45 seconds) with only one ordering of segments rather than two.

#### Preprocessing

Our preprocessing pipeline was similar to prior studies^7,73^. Electrode responses were common-average referenced to the grand mean across electrodes from each subject. We excluded noisy electrodes from the common-average reference by detecting anomalies in the 60 Hz power band (measured using an IIR resonance filter with a 3dB down bandwidth of 0.6 Hz; implemented using MATLAB’s iirpeak.m). Specifically, we excluded electrodes whose 60 Hz power exceeded 5 standard deviations of the median across electrodes. Because the standard deviation is itself sensitive to outliers, we estimated the standard deviation using the central 20% of samples, which are unlikely to be influenced by outliers (we divided the range of the central 20% of samples by that which would be expected from a Gaussian of unit variance). After common-average referencing, we used a notch filter to remove harmonics and fractional multiples of the 60 Hz noise (60, 90, 120, 180; using an IIR notch filter with a 3dB down bandwidth of 1 Hz; the filter was applied forward and backward; implemented using MATLAB’s iirnotch.m).

We measured integration windows from the broadband gamma power response timecourse of each electrode. We computed broadband gamma power by measuring the envelope of the preprocessed signal filtered between 70 and 140 Hz (implemented using a 6^th^ order Butterworth filter with 3dB down cutoffs of 70 and 140 Hz; the filter was applied forward and backward; envelopes were measured using the absolute value of the analytic signal, computed using the Hilbert transform; implemented using fdesign.bandpass in MATLAB). We have previously shown that the filter does not strongly bias the measured integration time because the integration window of the filter (∼19 milliseconds) is small relative to the integration window of the measured cortical responses^7^. Envelopes were downsampled to 100 Hz. We detected occasional artifactual time points as time points that exceeded 5 times the 90^th^ percentile value for each electrode (across all time points for that electrode), and we interpolated these outlier time points from nearby non-outlier time points (using “piecewise cubic Hermite interpolation” as implemented by MATLAB’s interp1.m function).

As is standard, we time-locked the iEEG recordings to the stimuli by either cross-correlating the audio with a recording of the audio collected synchronously with the iEEG data or by detecting a series of pulses at the start of each stimulus that were recorded synchronously with the iEEG data. We used the stereo jack on the experimental laptop to either send two copies of the audio or to send audio and pulses on separate channels. The audio on one channel was used to play sounds to subjects, and the audio/pulses on the other were sent to the recording rig. Sounds were played through either a Bose Soundlink Mini II speaker (at CUMC) or an Anker Soundcore speaker (at NYU). Responses were converted to units of percent signal change relative to silence by subtracting and then dividing the response of each electrode by the average response during the 500 ms before each stimulus.

We selected electrodes with a significant test-retest correlation (Pearson correlation) across the two presentations of each stimulus. Significance was measured with a permutation test, where we randomized the mapping between stimuli across repeated presentations and recomputed the correlation (using 1000 permutations). We used a Gaussian fit to the distribution of permuted correlation coefficients to compute small p-values^74^. Only electrodes with a highly significant correlation relative to the null were kept (*p <* 10^−5^). We also excluded electrodes where the test-retest correlation fell below 0.05.

Following standard practice, we localized electrodes as bright spots on a post-operative computer tomography (CT) image or dark spots on a magnetic resonance image (MRI), depending on which was available. The post-op CT or MRI was aligned to a high-resolution, pre-operative MRI that was undistorted by electrodes. Each electrode was projected onto the cortical surface computed by Freesurfer from the pre-op MRI, excluding electrodes greater than 10 mm from the surface. We used the same correction procedure as in our prior studies to correct gross-scale errors by encouraging points that are nearby in 3D space but far apart on the 2D cortical surface (e.g., two abutting gyri) to preferentially be localized to regions where sound responses are common.

#### Estimating integration windows from noisy neural responses

Neural responses are noisy and thus never produce the same response even if the response is context invariant. Moreover, we are limited in the number of segment durations and segments that we can test due to limited experimental time with patients. To address these challenges, we measure a noise ceiling for our cross-context correlation when the context is identical using repeated presentations of the same segment sequence. We then estimate the integration window that best predicts the cross-context correlation by finding a parametric window (Gamma distributed) that best predicts the cross-context correlation pooling across all lags and segment durations. The predictions are computed by multiplying a noise-free prediction from the parametric model window by the measured noise ceiling. We have previously described and justified our parametric model in detail and have extensively tested the method, showing that it can correctly estimate integration windows from a variety of ground truth models without substantial bias using noisy, broadband gamma responses with similar signal-to-noise ratios as those in actual neural data^7^.

In our original formulation, the Gamma-distributed window was parametrized by three parameters that control the width, delay, and shape of the window. We found previously that the best-fitting delay is highly correlated with the width and is close to the minimum possible value for a given width and shape, and therefore constrained the delay to take this minimum value to reduce the number of free parameters. For a structure-yoked response, we predict that the window will scale with the temporal scaling of the stimulus structures, which will change the width but not the shape. We therefore constrained the shape of the window to be the same for compressed and stretched speech. Results were similar, but with higher variance when the shape was unconstrained, likely in part due to an increased number of free parameters (**Fig S2**). We also measured the average cross-context correlation within each annular ROI as a simple, model-free way of assessing the average integration window for stretched and compressed speech (**Fig 3E**). The results of this model-free analysis support our model-dependent results by showing that integration windows increase in non-primary regions but change little with structure duration.

We also applied our parametric window analysis to our STRF and phoneme integration models which revealed the expected result with time-yoked integration for our STRF model and structure-yoked integration for our phoneme model (**Fig S3**). It was not possible to apply our parametric window analysis to our DNN model because unlike the brain the model’s responses are acausal and a Gamma-distributed window is thus inappropriate.

#### Timecourse rescaling

We measured the degree to which the neural response time course rescaled by correlating the response time course to stretched and compressed speech after rescaling the time course for the compressed speech (accomplished via resampling, i.e. upsampling by a factor of 3, and then discarding the additional time points). After rescaling, we measured the average correlation between all pairs of stimulus repetitions, and we measured a ceiling for this across-condition correlation (stretched vs. rescaled compressed) by measuring the average test-retest correlation across repetitions for the same condition (stretched vs. stretched and rescaled compressed vs. rescaled compressed). We only used responses to the intact 27-second stimuli for this analysis. We applied the same analysis to the trained and untrained DNN models.

#### Statistics

Unlike most intracranial studies, we had responses from enough participants to perform across-subject statistics, and when possible used a linear mixed effects model to account for subjects as a random effect (using lmefit.m in MATLAB). To evaluate whether the integration windows differed between stretched and compressed speech we used the following model:

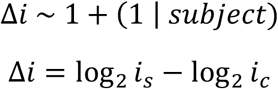

which models the logarithmic difference in integration windows between stretched (*i*_*s*_) and compressed (*i*_*c*_) speech using a fixed effects intercept plus a subject-specific random intercept. We then tested whether the fixed effects intercept differed significantly from 0 using an F-test (as implemented by coefTest.m with Satterthwaite approximation used to estimate the degrees of freedom; a full covariance matrix was fit in all cases).

To examine the effect of distance (*d*) on overall integration windows, we modeled the mean integration window for stretched and compressed speech as a function of distance to primary auditory cortex:

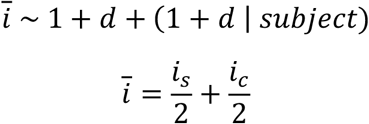

To evaluate the effect of distance on structure yoking, we modeled the difference in integration windows (which is proportional to the structure yoking index) as a function of distance:

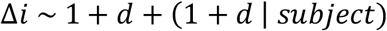

To evaluate whether the structure yoking increased at longer timescales, we modeled the difference in integration windows as a function of the overall integration window:

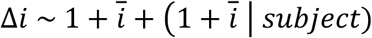

We evaluated whether there was a significant effect of timecourse rescaling by fitting a model on the difference in correlations between rescaled and non-rescaled responses:

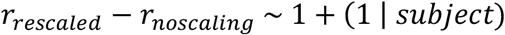

Bootstrapping was used to compute error bars (**Fig 3C&D**), resampling both subjects and electrodes with replacement (resampling subjects and for each subject resampling electrodes). Error bars plot the central 68% interval (equivalent to one s.d.) of the bootstrapped distribution.

## Acknowledgments

We thank Laura Long for help with data collection. This study was supported by the National Institutes of Health (NIDCD-K99-DC018051, NIDCD-R00-DC018051 to S.V.N.-H.). The funders had no role in study design, data collection and analysis, decision to publish or preparation of the manuscript.

## Author contributions

S.V.N.-H. and M.K. collected data for the experiments described in this manuscript. O.D., W.D., G.M.K. and C.A.S. collectively planned, coordinated and executed the neurosurgical electrode implantation needed for intracranial monitoring. S.V.N.-H. performed the analyses of intracranial data and M.K. performed the analyses of the phoneme/STRF and DNN models. S.V.N.-H. designed and implemented the experiment with mentorship and guidance from N.M. and A.K. S.V.N.-H. wrote the paper with feedback from N.M. and A.K., as well as all other co-authors.

## Supplemental Figures

**Supplemental Figure 1.**
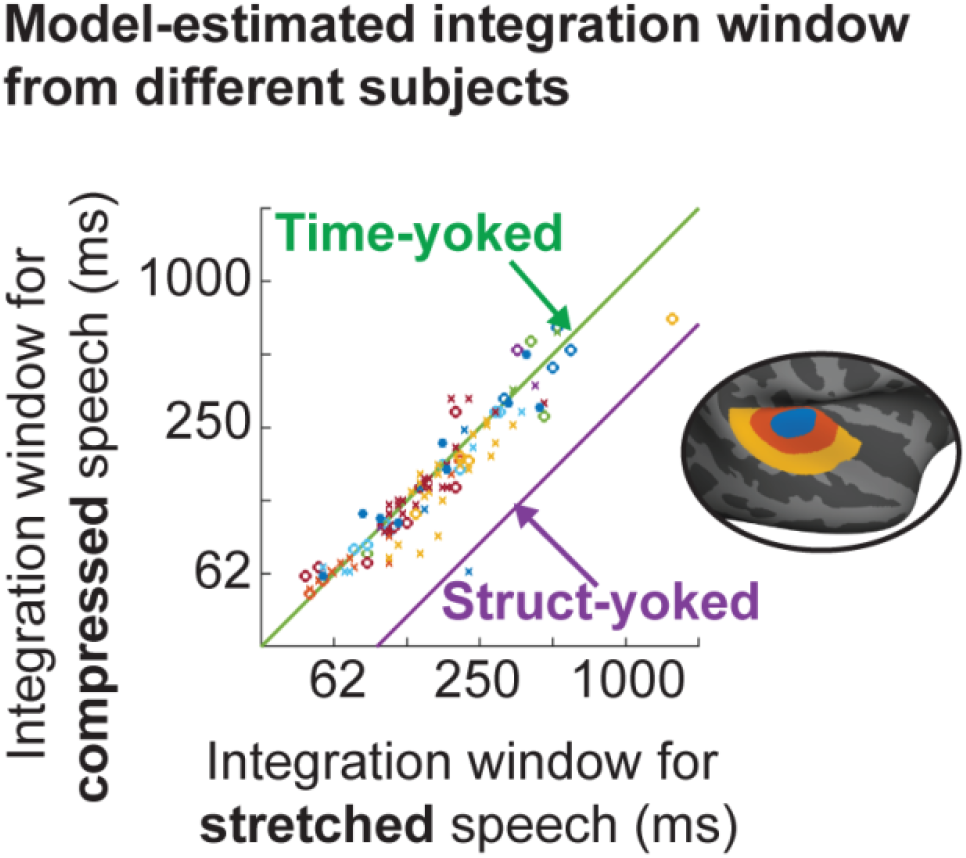
Consistency across participants. Integration windows for stretched (x-axis) and compressed (y-axis) speech for all sound-responsive electrodes. Electrodes from the same participant are given the same symbol/color. The format is otherwise the same as in Figure 3B.

**Supplemental Figure 2.**
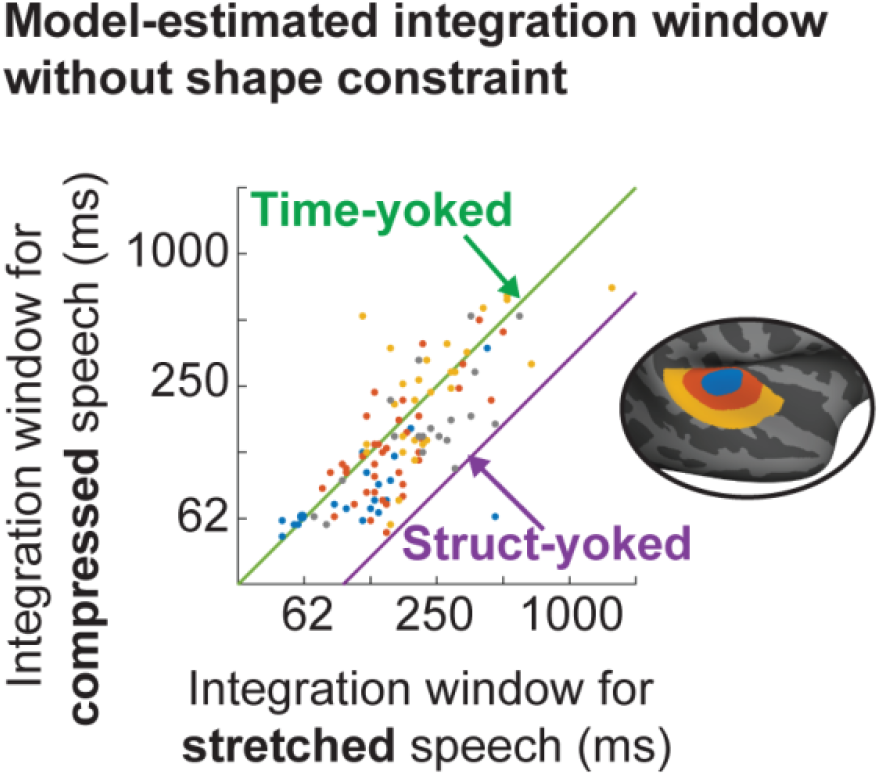
Unconstrained window shape. To reduce the number of free parameters and increase robustness, we constrained the shape of the Gamma-distributed window to be the same for stretched and compressed speech. This figure shows the results when the shape is unconstrained. Results are similar but the variance is higher, as expected. The format is the same as in Figure 3B.

**Supplemental Figure 3.**
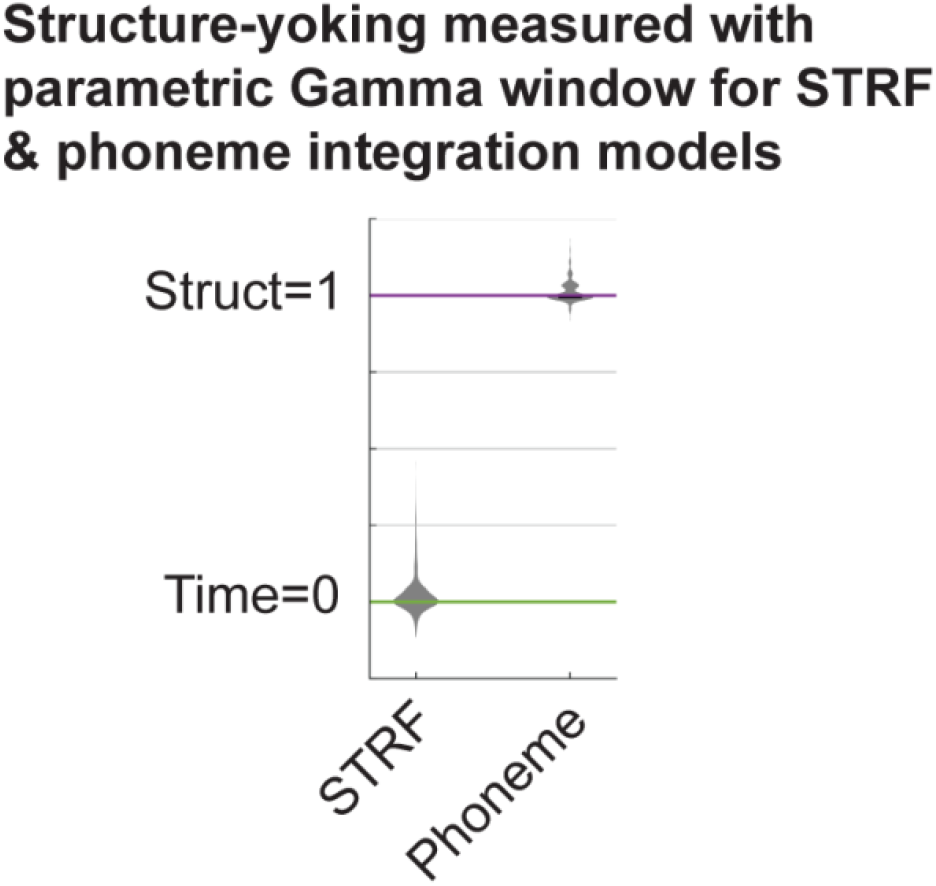
Structure yoking for phoneme and STRF models estimated using parametric Gamma window. We used a simple, model-free approach to estimate integration windows from STRF and phoneme integration models since the model responses are noise-free and not constrained by limited data. Here, we show that structure-yoking values are similar when estimated using the parametric, Gamma-distributed window applied to neural data. The format is the same as in Figure 2C.

